# A tale of two distinct actin networks underlies the entire life cycle of focal adhesion

**DOI:** 10.1101/2025.11.12.688129

**Authors:** Rico Dong, Karan Ishii, Sergey Plotnikov, Jian Liu

**Author notes:** Corresponding Authors (S.P.); (J.L.). Equal Contributions (Rico Dong, Karan Ishii).

## Abstract

Cell assembles focal adhesion (FA) to transmit the stress fiber (SF)-based actomyosin contraction onto the extracellular matrix (ECM) for mesenchymal migration, essential for many physiological processes (*e.g.*, development and wound healing). To transmit force efficiently, both FA and SF contractility are built as “clutches” and in positive feedback with each other; conversely, the SF-engaging FA imposes a strong cell-ECM anchorage and must be disassembled timely to facilitate the cell migration. How the cell balances the two opposing roles of FA in cell migration is an open question. Particularly, it is not well-understood how a cell builds the FA *de novo* to clutch with SF and disassemble the clutches when needed in a coherent manner. Combining theory and experiments, we show that the entire life cycle of FA is seamlessly orchestrated by the FA-localized spatiotemporal coordination between retrograde actin flow and SF, without destroying the FA constituent molecules. Retrograde actin flow drives the centripetal growth of nascent FA, paving the way for SF engagement. The SF further stabilizes the growing FA into maturation via the positive feedback that clutches the contractility with the FA. Finally, the retrograde actin flow increase, in coupling to the local cell edge retraction, tugs the mature FA in the proximal direction that relaxes the associated SF contractility, turns off the clutching, and triggers the FA disassembly. Our finding sheds light on the organizational principles that cell streamlines the mechanochemical interplay between FA, actin cytoskeleton, and cell edge dynamics for efficient cell migration.

## Introduction

Cell assembles focal adhesion (FA) *de novo*, an integrin-based transmembrane linkage with the extracellular matrix (ECM), through which it exerts the traction forces to power the mesenchymal migration (1–4). As a primary mechanosensitive organelle of the cell, FA adapts the cell’s traction force according to the mechanics of the local ECM to mediate directed cell migrations (*e.g.*, durotaxis (5–7)), which are crucial for many physiological processes such as development (8, 9), tissue formation (10, 11), and cancer metastasis (12–14). As a strong cell-ECM anchorage, the FA must be disassembled in a timely manner to facilitate cell migration (15, 16). Despite its essential role in cell physiology and the intensive research, however, the coherent physical mechanisms underlying the entire life cycle of a FA are not well understood.

FA has a layered structure (17–19); in a simplified picture, integrin molecules bind to ECM on one end and engage with the actin cytoskeleton via FA adaptor proteins (*e.g.*, talin, vinculin, and paxillin) on the other end. A FA starts its life journey as a small nascent FA (∼ 100-200 nm in diameter) near the cell edge, where cell spawns approximately 10s -100s of nascent FAs (1, 2). While many of these nascent FAs turn over rapidly, some manage to grow centripetally by interacting with the retrograde actin flow, which results from a combination of branched actin network polymerization and cell edge protrusion. Some of these directionally growing FAs then mature into micron-sized transmembrane domains and engage with stress fibers (SF), which mainly consist of actomyosin filaments (1, 2). The mature FA transmits SF-mediated contraction onto the ECM that drives cell migration. As the cell moves forward, the mature FA – while remaining largely stationary with respect to the ECM – “slides” backward relative to the cell (1, 2). When these mature FAs reach the cell rear end, they must disassemble timely (15, 16); otherwise, they will hinder cell migration and may even rip the migrating cell leaving some integrins behind (20).

Hereby, the cell faces a design problem is: To facilitate efficient force transmission, both mature FAs and SF-mediated actomyosin contractility are built as a “clutch”: the FA strengthens its ECM linkage in response to traction force (21), and the contractility is potentiated upon resistance (22) (*aka* catch bond-like dynamics). As such, a mature FA and its engaging actomyosin contractility are in clutching by positively feeding back with each other (23). This precipitates the following pressing questions: How can the cell turn off this positive feedback to disassemble the clutching FA? How does the mechanism of mature FA disassembly play out in the coherent process of FA’s entire life cycle, beginning from its infancy, to maturation, and eventually to demise?

One simple answer is that the cell can disengage actomyosin contractility to induce the mature FA disassembly. Indeed, experiments suggest that calcium influx activates the calpain-mediated pathway, which cleaves FA adaptor and signaling proteins (*e.g.*, talin and FAK) and severs the linkage between the integrin and actin cytoskeleton, thereby triggering FA disassembly (24, 25). However, if this calcium-mediated cleavage is the main mechanism of mature FA disassembly, then it would have to solve two control problems. First, given that calcium is highly diffusive and calcium channels do not exclusively localize at mature FAs, how does the cell distinguish the mature FAs that need to be disassembled from those don’t? Second, how does the calpain-mediated pathway of mature FA disassembly align with the cost-effective strategy that any cell has to consider in the long run? In this case, the cell migration would entail many talin molecules to be cleaved from the mature FAs and meanwhile, many new talin molecules to be synthesized for assembling new FAs. Since both protein synthesis and cleavage cost energy and building material for the cell, they would put quite a burden on the cell physiology and metabolism. It is not a question of whether the cell could exploit this calpain-mediated pathway for mature FA disassembly, as experiments clearly demonstrate the possibility (24, 25). Rather, the real question is whether the primary mechanism of mature FA disassembly hinges on non-cleavage pathways so that the building materials for FA can be readily re-used and energy can be saved.

Indeed, nascent FAs turn over without invoking the calpain-mediated pathway. This begs the fundamental and unanswered questions: Can the same mechanism underlie both FA assembly and disassembly by just changing the operating parameters, instead of invoking additional pathways? In other words, how does FA bear its own demise in the “code” of assembly?

Combining theoretical modeling and experimental testing, we present evidence that the entire life cycle of FA is coherently orchestrated by spatiotemporal coordination between the retrograde actin flow and SF, without the need to invoke the calcium-calpain-cleavage pathway. Specifically, retrograde actin flow promotes the centripetal growth of nascent FA by interacting with FA catch-slip bond dynamics via a reaction-diffusion-convection process. This results in a spatial gradient of FA-localized PTK activity tapering toward the proximal end of the growing FA, which sets the stage for FA-localized PTP activation and SF engagement. The SF further stabilizes the growing FA into maturation via the positive feedback between the catch-bond dynamics of the actomyosin contractility and the FA. Finally, the increase in the retrograde actin flow, in coupling to the local cell edge retraction, tugs the mature FA in the proximal direction; this relaxes the associated actomyosin contractility, turns off the clutching, and triggers the FA disassembly. Our finding not only provides a unified mechanism that the FA-localized crosstalk between retrograde actin flow and SF underlies the entire life cycle of FA from the initiation, growth, maturation, and to turnover, but also sheds light on the organizational principles that cell exploits to streamline the mechanochemical interplay between cell edge dynamics, actin cytoskeleton, and FA for efficient cell migration.

## Results

### Model Description

To dissect the fundamentals of FA’s life cycle, we aimed to construct the theoretical model that coherently captures how a single FA grows from a nascent state in a lamellipodium (LP) to a mature state in lamellum (LM), and finally, to disassembly of the mature FA at the rear of the cell (Figure 1A). Expanding upon our established work (7), the essence of our model is that FA-localized mechanochemical interplays between retrograde actin flow, SF, and the FA catch-slip bond dynamics orchestrates not only the FA growth, but the entire progression of FA development from birth to death. Importantly, key elements of these mechanochemical interplays were supported and characterized by *in vitro* and *in vivo* experiments and therefore strongly constrain the theoretical model. The theoretical model organically synthesizes these ingredients into a coherent picture of FA deveopment on a functional module level as follows.

**Figure 1:**
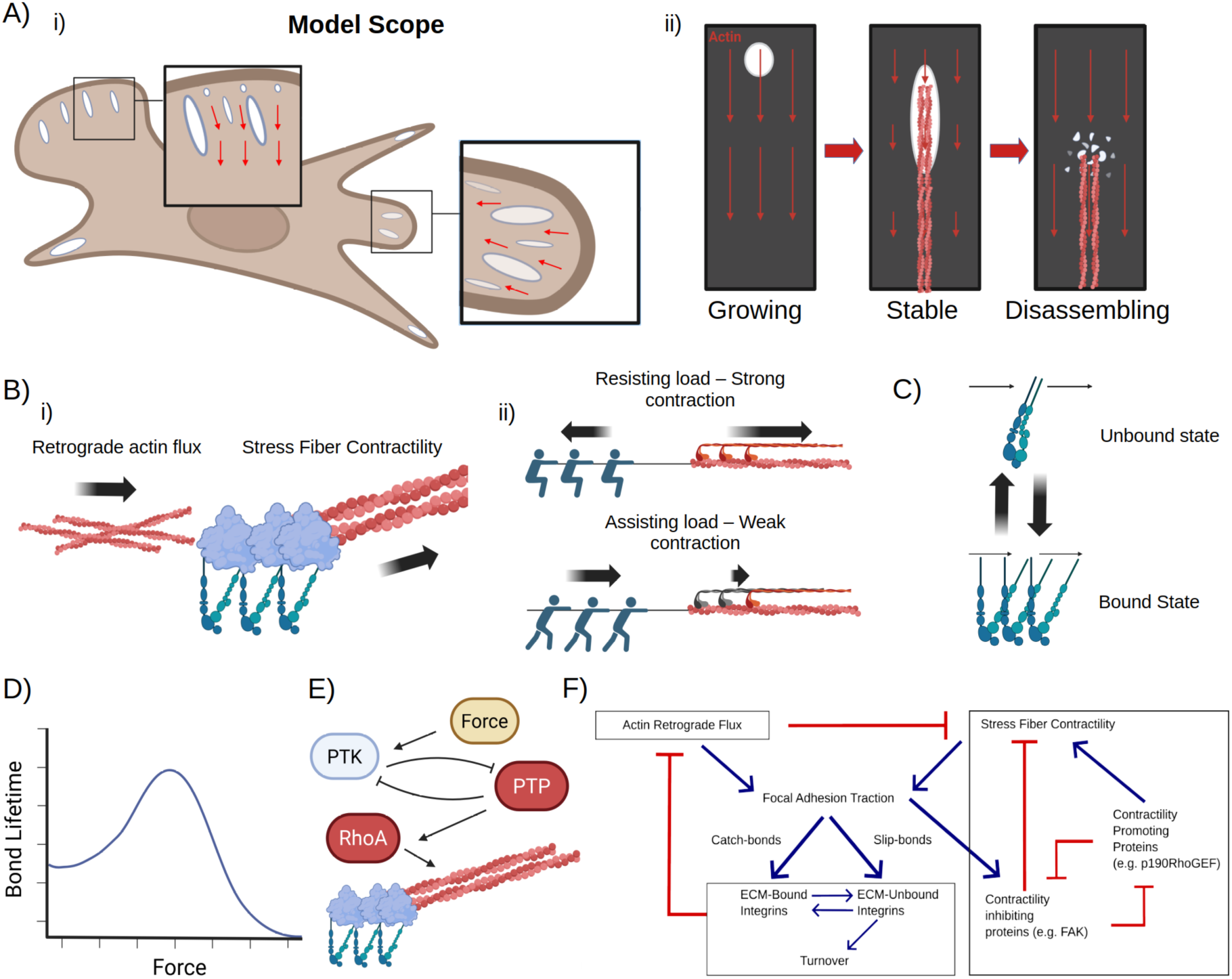
Model schematic. (A) The model scope encompasses the complete lifecycle of FAs (Ai, Aii), from their initial assembly, to their maturation and stabilization, up to their disassembly. (B-E) Outlines of key modules. (Bi) Actin module. Both retrograde actin flux and stress fiber contractility exert force at FAs through interactions with FA components. (Bii) Load-dependence of nonmuscle myosin II. Resisting loads drive strong contraction, while assisting loads weaken contraction. (C) Two-state model of FA components. Components can be either bound (mimicking ECM-Integrin-Adapter complexes) or unbound (mimicking all other states of FA components). (D) Catch-slip bond behavior of FA components. At intermediate forces, FA components are reinforced, while at high or low forces, FA components disassemble. (E) PTK-PTP signaling module. PTP and PTK are assumed to be mutually inhibitory; PTK is activated by force, while PTP promotes stress fiber formation and contractility through the RhoA pathway. (F) Wiring diagram summarizing the model.

#### 1. Actin module

This module includes retrograde actin flow and SF that both exert traction forces on FA. For the sake of simplicity, we imposed a fixed rate of retrograde actin flow at the cell edge, without explicit describing the cell edge protrusion/retraction dynamics (Figure 1B(i)). The retrograde actin flow mechanically tug the FA due to the binding affinity between the FA adaptor proteins and the actin (26). Additionally, SF-mediated actomyosin contractility itself is load dependent (22): Resisting loads stimulate stronger myosin II contraction, whereas facilitating loads weaken it (Figure 1B(ii)). It is the contractilty that positively feeds back with the FA clutching for efficient force transmission (see below).

#### 2. FA structural module

We simplified FA structural components into two inter-convertible states (Figure 1C): 1) bound complex that depicts the ECM-integrin-adaptor linkage and is the main force-bearing FA component, and 2) unbound complex reflecting the collective behavior of integrin-adaptor complex, integrin, and adaptor, which disengage from the ECM. When the bound complex converts to the unbound (*i.e.*, the ECM-integrin-adaptor complex becomes ECM and integrin-adaptor, integrin, or adaptor itself), the local retrograde actin flow drifts the unbound complex downstream (27). The unbound complexes can either convert to the bound state (*aka* rebind to ECM) or turn over into cytosol where they become well-mixed with a fixed intracellular concentration.

#### 3. FA catch-slip bond dynamics

In a high-force limit, the FA bound state like any chemical bond behaves as a slip bond. That is, when the tugging force is too high, it will break the bonds in the ECM-integrin-adaptor linkage, converting the bound state to unbound state (Figure 1D). Crucually, in the low-force regime, the FA bound state behaves as a catch bond (*aka* clutch (21)) (Figure 1D): *i.e.*, upon traction force from the retrograde actin flow-mediated tugging or SF-mediated contraction, more unbound complexes convert to the bound complex, which is based on experiments showing FA adaptors (*e.g.*, talin) under stretch expose more of the cryptic binding site to recruit more FA adaptor proteins (*e.g.*, vinculin) and integrin (28). This way, the FA strengthens its mechanical linkage with ECM upon traction force, like a clutch.

#### 4. FA-localized PTK/PTP signaling module

PTK (*i.e.*, the Src-FAK-CAS functional module) localizes to FA and its kinase activty is mechanosensitive (29). This is based on the experiments showing that the greater the local traction force, the more its kinase activity increases (Figure 1E) (29). Importantly, PTK is in mutual antagonism with PTP (30, 31), which is upstream of RhoA-mediated SF formation (Figure 1E) (30–33).

To mathematically describe the model, we used a system of coupled partial differential equations (PDEs) to capture the reaction-diffusion-convection process of FA development, which hinges on the mechanochemical feedback between the functional modules (Figure 1F). As the height scale around FAs is significantly smaller than its length and width scales (∼200 nm height in the lamellipodium (LP), ∼20 µm cell length/width), we approximated the system as 2D. Our simulation focuses on the life cycle of a single FA and initializes in a wide patch of LP spanning 5.5 µm long by 8 µm wide. From there, we fixed the retrograde actin flow velocity at the leading edge and simulated actin depolymerization at a fixed distance from the cell edge, representing the LP/LM interface (34). To describe that the FA changes its relative location inside the cell due to the cell migration (*e.g.*, deep inside the LA where the retrograde actin flow vanishes), we then modified these simulation conditions accordingly. Details in the mathematical formulation and numerical simulations are relegated to **SI**.

To capture the physical mechanism underlying the entire life cycle of a FA, we focused on three key developmental stages of FA with the theoretical model as the coherent mechanistic underpinning. Below, we dissected first how a nascent FA grows, followed by how a growing FA gets stabilized and matured, and lastly, how a mature FA disassembles. Importantly, these theoretical endeavors not only help explain novel experimental observations in previous publications but predict nontrivial FA dynamics that were supported by our own experiments in this paper. Together, our findings piece multiple lines of evidence into a coherent picture of FA’s life cycle, driven by the spatiotemporal coordination between the branching actin network and stress fiber.

### Actin retrograde flow engages with FA catch-slip bonds to drive FA centripetal growth

To dissect how a nascent FA grows, we focused on the simplest case. We fixed the retrograde actin flux velocity at the leading edge and imposed open boundary conditions everywhere else. We then initialized a FA with a diameter of 200 nm containing only FA anchored components and PTP/PTK in chemical balance (Figure 2A). In a nutshell, our model captures the essential features of the observed directional FA growth (35), predicts that mechanochemical interactions between retrograde actin flux and FA catch-slip bond behaviors play a critical role in this process (Figures 2 and 3), and explains the novel observations of the mechanochemical requirements for initial cell adhesion (Figures 2 and 3).

**Figure 2:**
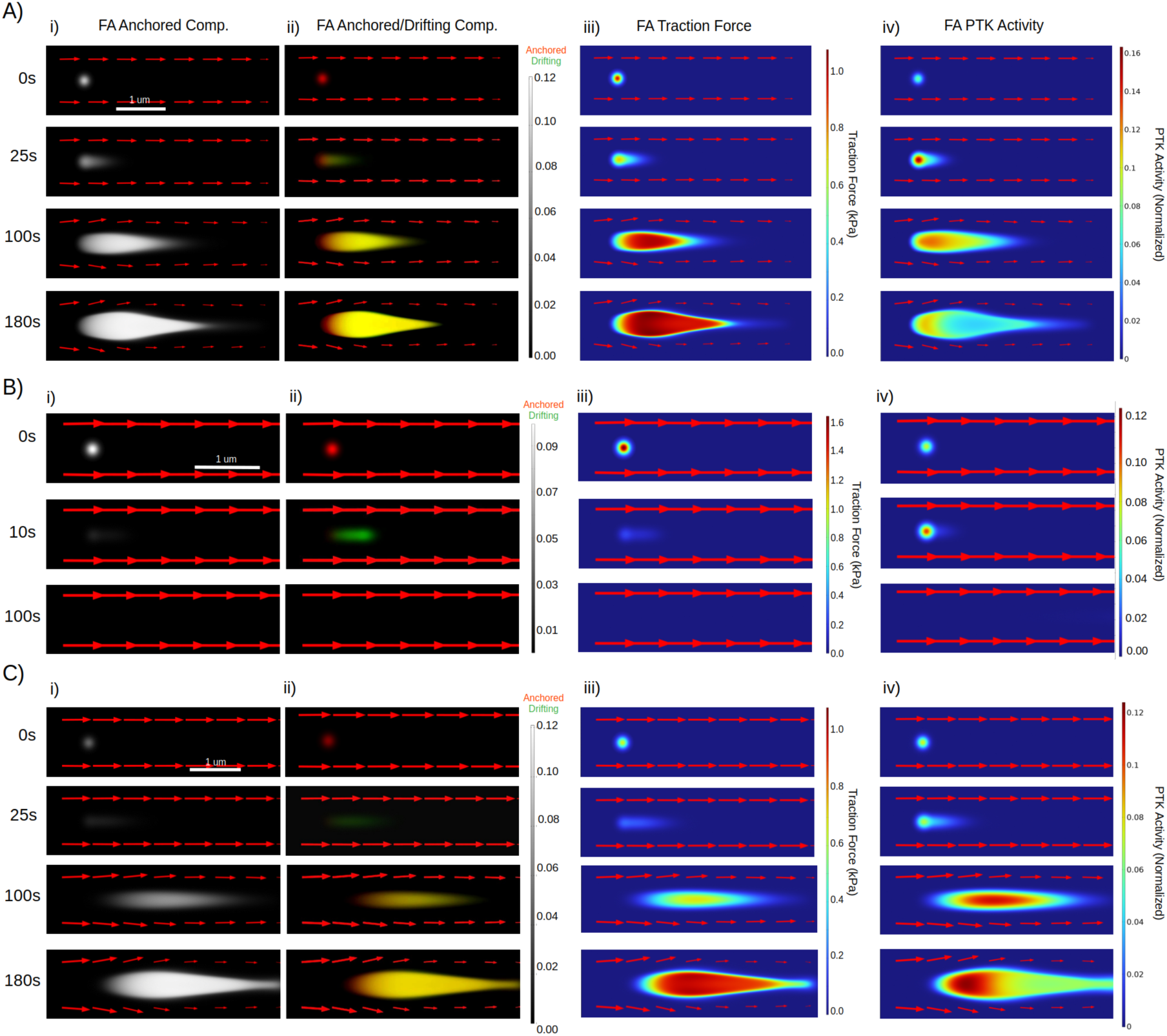
Nascent adhesions undergo centripetal growth, sliding or washout in response to actin flow. (A, B, C): Predicted spatiotemporal profile of focal adhesion anchored components, unanchored components, PTK activity and traction force at different parameter regimes. Arrows indicate retrograde actin flux; length of arrows correlate with flux magnitude. (A) Centripetal FA growth. Flow rate = 40nm/s, initial FA Density = 0.1. (B) Focal adhesion washout. Flow rate = 70nm/s, initial FA Density = 0.1. (C) Focal adhesion slipping, followed by growth. Flow rate = 52.5nm/s, initial FA Density = 0.05. (Ai, Bi, Ci): Spatiotemporal profile of anchored FA components over time. (Aii, Bii, Cii): Anchored and unanchored FA components over time. Unanchored components are in green, while anchored components are in red. (Aiii ,Biii, Civ): FA traction force in kPa over time. Red indicates higher traction forces, while blue indicates lower traction forces. (Aiv, Biv, Civ): Spatiotemporal profile of PTK activity over time. As the FA grows, it slows down the flow at the rear, which concentrates PTK activity at the front due to force-dependence of PTK. Red indicates higher PTK activity, while blue indicates lower activity.

**Figure 3:**
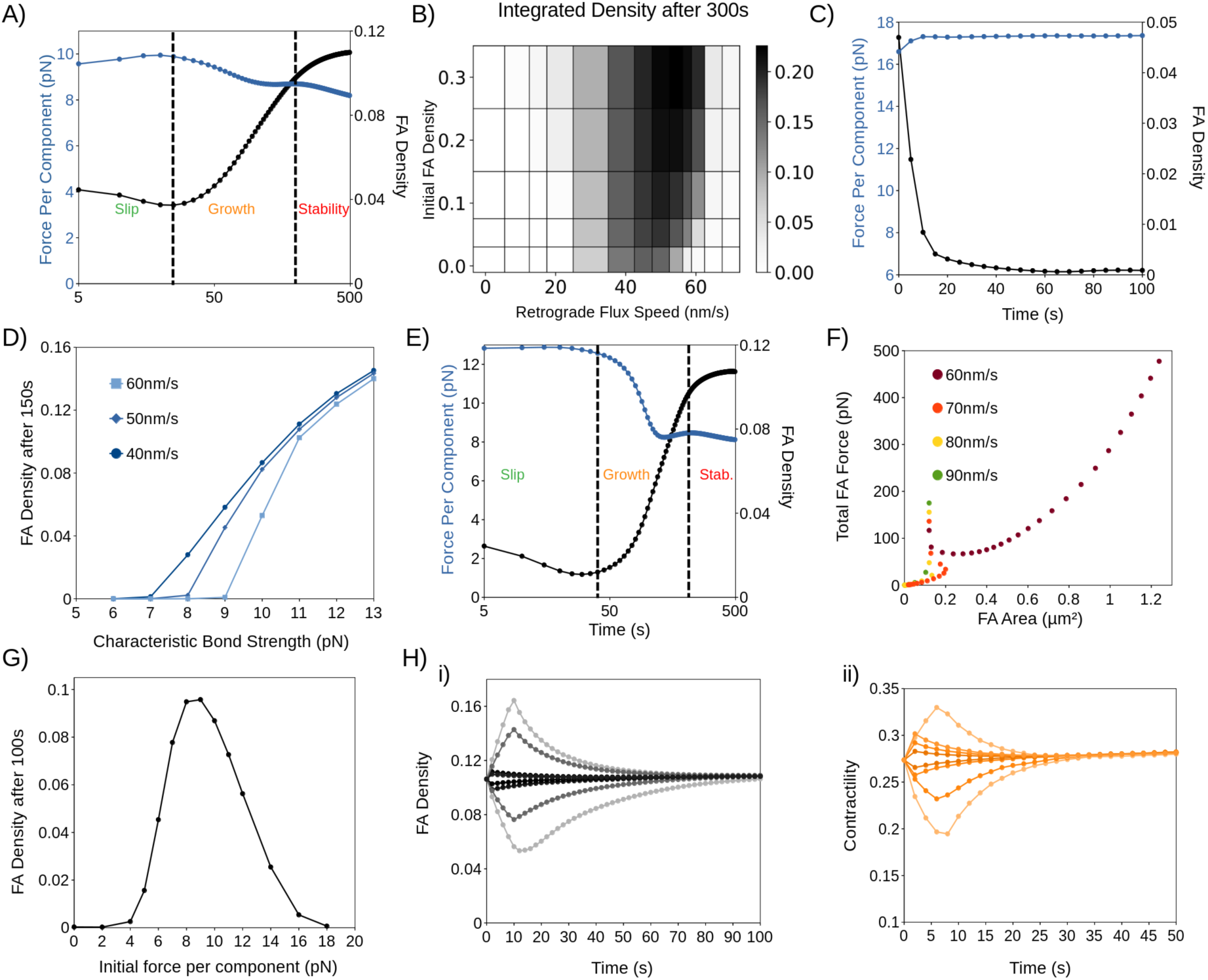
Interactions between actin flow and catch-slip bonds govern FA dynamics. (A) Average force per FA component and FA Density during centripetal growth (Figure 2A). From this point forwards, FA Density refers to average density of the anchored phase. Blue indicates force per component, while black indicates average FA density. B) Phase plot showing the dependence of FA growth (shading represents integrated FA density (units of µm^2), equal to average density multiplied by area) on retrograde actin flux speed and initial FA density. (C) Average force per FA component and FA Density during washout (Figure 2B). (D) FA Density after 150s at varying slip bond threshold forces, done at a fixed flow speed (40nm/s, 50nm/s, 60nm/s). (E) Average force per FA component and FA Density during slipping (Figure 2C). (F) Total force per FA with respect to FA area at different retrograde flow speeds (60nm/s, 70nm/s, 80nm/s, 90nm/s). (G) Plot depicting the average FA Density after 100s at a fixed force per component. (Hi, ii) FA Density (Hi) and Contractility (Hii) after perturbations from steady state with varying length (1-10s) and amplitude (0.05-0.1 1/s).

In the canonical case where a nascent FA grows directionally, the model suggests that actin retrograde flow invokes three effects in parallel that play out at each location within the FA (Figure 2A). First, the convection will drift unbounded FA components downstream. Consequently, this will shift the local chemical equilibrium so that more FA anchored components will become unbounded and enter the drifting population. This way, the actin retrograde flow weakens the FA (the slip-phase in Figure 3A). Second, the mechanical engagement with actin retrograde flow generates the traction force on the FA anchored components; this invokes the catch bond behavior of FA, assembling more FA anchored components into the FA. Consequently, as the FA is strengthened and makes the stronghold, it will slow down the local actin retrograde flow, attenuating the drifting effect. Third, while drifting with the actin retrograde flow downstream, some of the FA unbounded components (*aka* FA building materials) will deposit and become anchored onto the substrate. The combination of the second and third effects contributes to the centripetal growth of the nascent FA (the growth-phase in Figure 3A).

The contributions of the three effects to whether and how a nascent FA grows hinge on the strength of FA-substrate binding, relative to the convection by actin retrograde flow (Figure 3B). Hereby, the strength of FA-substrate binding includes a) the anchored component density of the nascent FA, b) how fast the FA unbounded component becomes anchored onto the substrate, and c) how fast the FA anchored components self-reinforce (catch bond) and convert into unbounded state (slip bond) upon the traction force, respectively. When the FA-substrate binding is strong (Figure 2A), the nascent FA will remain anchored onto the substrate at its distal end (facing the cell edge), while undergoing the centripetal growth at its proximal end (facing the cell interior). Conversely, when the strength of FA-substrate binding is weak compared to the convection effects, the nascent FA will be washed away by actin retrograde flow (Figures 2B and 3C), which is the fate of many nascent FAs in the initial cell adhesion processes (1, 2).

To further dissect the roles of catch-slip bonds in FA growth, we leveraged the model by focusing on the threshold traction force per FA anchored component (Figure 3D), beyond which the anchored->unbounded state transition rate increases drastically. This threshold force is characteristic of slip bonds and defines the FA-substrate bond strength. The model predicts that for a given rate of actin retrograde flow, there exists a threshold FA-substrate bond strength, only above which, can a nascent FA undergo centripetal growth; otherwise, it will dissolve upon engaging with actin retrograde flow (Figure 3D). Furthermore, this threshold bond force is predicted to scale with the rate of actin retrograde flow. This lends an explanation for the novel observation that the initial cell adhesion, independent of actomyosin contractility, requires a threshold integrin-substrate bond strength that scales with the effective membrane tension (36). According to our previous work (37, 38), the larger the membrane/cortical tension, the harder it is for actin polymerization to advance the cell edge, resulting in a faster actin retrograde flow.

Interestingly, when the strength of FA-substrate binding is around the transition regime - for instance, when the nascent FA has a relatively low density of anchored components - the model predicts a novel phenomenon: Initially, the nascent FA cannot hold onto the substrate at its distal end; instead, the convection by the actin retrograde flow is predominant (Figures 2C and 3E). During this process, the entire nascent FA undergoes repetitive episodes of unbinding-convection-anchoring (Figure 2C). In other words, the nascent FA “treadmills” downstream. Eventually, the unbounded FA components anchor onto the substrate, and the system moves towards a parameter regime that permits strong FA-substrate binding. From this point onward, the FA makes a stronghold at its distal end while growing centripetally at its proximal end. Consequently, the slip phase of nascent FA growth is much longer than the nominal case (compare the Figures 3B and 3E). This provides an explanation for the observation that nascent FAs can “slide” a long distance before growing into mature FAs (39). Critically, it is our prediction that FA “sliding” is really the adhesion treadmilling with the actin retrograde flow, rather than the entire FA moving as an intact physical object.

The model so far suggests that centripetal FA growth arises from the mechanochemical interplay between FA catch-slip behavior and actin retrograde flow. How does a growing FA stabilize and mature? What role does centripetal growth play in FA maturation?

### Stress fiber contractility self-reinforces with FA catch bonds to promote FA maturation

Our model predicts that 1) the reaction-diffusion-convection process of FA centripetal growth creates a FA-localized traction gradient that decreases toward the FA proximal end (Figure 2A(iii)), and 2) this mechanical gradient paves the way for the SF engagement by modulating the FA-localized antagonism between PTP and mechanosensitive PTK activity (Figure 2A(iv)).

Specifically, active PTK is recruited early on in the nascent FA as evidenced in experiments (*e.g.*, (40)) and implemented as an initial condition in the model. Because PTK and PTP are mutually antagonistic (30, 31), the chemical balance between them is in flux as they compete, with the PTK being predominant over PTP in nascent FAs. As the nascent FA grows centripetally, the anchored FA components put up a stronghold by tugging on the retrograde actin flow, which results in a high traction near the FA distal end. Conversely, the unbound FA components drift with the retrograde actin flow downstream and accumulate toward the FA proximal end, transmitting a lower traction onto the ECM. Because PTK activity is potentiated by the mechanical forces (29), it continues to dominate over PTP near the FA distal end. In contrast, the low traction near the FA proximal end tips the PTK/PTP balance in favor of PTP activation. Since PTP is a positive regulator of actomyosin contractility, this leads to SF engagement at the rear of the centripetally growing FA (Figure 2A(iii)). As the FA grows larger, it impedes the retrograde actin flow (41), which contributes even less to the local traction force near the FA proximal end. Meanwhile, the SF-mediated actomyosin contractility becomes the dominant source of traction force in the proximal half of the growing FA. This promotes the stabilization and maturation of the FA (the stability phase in Figure 3A). Finally, as the cell edge protrudes forward, the FA that was growing in LP now ends in LA, where the retrograde actin flow diminishes and so does its engagement with the FA distal end. As a result, the FA-localized PTK activity yields to PTP, allowing the adhesion to fully engage with the SF and mature.

The above picture indicates that the FA-localized traction has two different origins from the branching actin network and SF, respectively, the relative contribution of which couples to the FA developmental process. In mature FAs, the spatial profile of force is maintained by actomyosin, the FA traction force is predicted to scale linearly with the FA area at a slope equal to the optimal traction force (Figure 3F). The exception to this is when the actin retrograde flux is significantly stronger than FA-substrate binding. In this case, the nascent FA will momentarily experience a very high force relative to its area before dissolution, which scales with the speed of actin flow as well as the initial density of the nascent adhesion (Figure 3H). When plotted alongside FAs undergoing centripetal growth, these nascent FAs appear as points with very high traction but very small area. This explains the observation of the “V”-shaped force-area relationship (42): *i.e.*, the traction force scales linearly with the FA area in mature FAs, whereas small/nascent FAs are under very high force.

This same mechanochemical picture implies that 1) FA-localized PTK activity near the FA distal end is high when the growing FA tugs with the retrograde actin flow, and 2) it decreases progressively as the FA matures in LA where the local retrograde actin flow vanishes. These mechanistic insight from our model provides a coherent explanation for the observations that the FA-localized PTK activity (*e.g.*, FAK) polarizes towards the FA distal end and becomes diminished in large FAs (43, 44).

We next interrogated the stability of a mature FA actively transmitting traction forces on the ECM. According to our model, FA stability and maturation are driven by positive feedback of catch bond-like dynamics between actomyosin contractility and FA. That is, the FA strengthens its linkage with ECM upon SF-mediated actomyosin contraction; and the contractility is potentiated by an increased resisting load (in the form of a strengthened FA-ECM linkage) and conversely, disengages upon facilitating load (22). To further understand the mechanistic role of this positive-feedback driven clutch in FA stability, we leveraged the model to examine how the steady-state FA density and traction forces respond to perturbations. The model predicts that actomyosin maintains the force per component at an intermediate range of mechanical force that maximizes catch bond engagement and minimizes slip bond breaking (Figure 3G). Importantly, this steady state is maintained against perturbations (Figure 3H). This is because, when the FA density was artificially decreased, the traction force per FA anchored component increases. This invokes FA catch-bond dynamics by enhancing the recruitment of more FA anchored components to this weakened FA, which strengthens the FA-ECM linkage and drives the system back to the steady-state. Likewise, when actomyosin contractility is artificially decreased, FA-localized PTP remains unperturbed in promoting SF contractility, restoring it back to the steady-state level. This way, the mature FA remains stable against perturbations and enables efficient force transmission.

However, this stability is a double-edged sword. Mature FAs must disassemble in a timely manner to facilitate cell migration. However, disassembling such a large stable structure is inherently a nontrivial task. Of course, substantial perturbations against the mature FA (*e.g.*, complete disruption of both branching actin networks and SF engaged to the FA, or loss of the entirety of actomyosin contractility) can lead to FA disassembly (Supplemental Figure S1). The real questions are: How are these drastic perturbations triggered inside the cell? Do they invoke additional pathways or exploit the same mechanism of FA growth and maturation as elaborated above?

### Calcium does not instigate mature FA disassembly

So far, our model suggests that mature FAs do not disassemble spontaneously nor upon moderate perturbations because of the self-stabilizing feedback loop between biochemical signaling and contractility (Figure 3H). What could trigger the mature FA disassembly?

Previous experiments demonstrated that calcium influx can trigger for FA disassembly through the activation of calpain-mediated cleavage of FA adaptor proteins (*e.g.*, talin and FAK) (24, 25). We therefore hypothesized that if calcium was indeed the primary trigger for mature FA disassembly, then we would expect some aspect of calcium dynamics to be clearly distinct between assembling and disassembling FAs.

To identify the distinction between assembling and disassembling FAs, we used a GCaMP6s calcium sensor paired to Paxillin-mCherry to visualize calcium specifically at FAs. Surprisingly, what we found was that FAs seem to assemble and disassemble independently of calcium (Figures 4A and 4B). Calcium flashes appear at both assembling and disassembling FAs, with little distinction in the frequency and amplitude of calcium flashes (Figures 4C and 4D). To ensure that cell-cell variation did not affect our results, we compared the FAs in the same cell by separating the amplitude of disassembling FAs from that of assembling FAs and computed the ratio between them (Figure 4E). Again, we observed only a 15% difference between assembling and disassembling FAs, with assembling adhesions exhibiting a higher calcium amplitude. Critically, there are numerous cases where calcium undergoes multiple flashes at a FA, yet little to no changes occur at the FA even after an extended period (*e.g.*, Figure 4A, B). This suggests that calcium is not the immediate instigator for the primary mechanism of mature FA disassembly.

**Figure 4:**
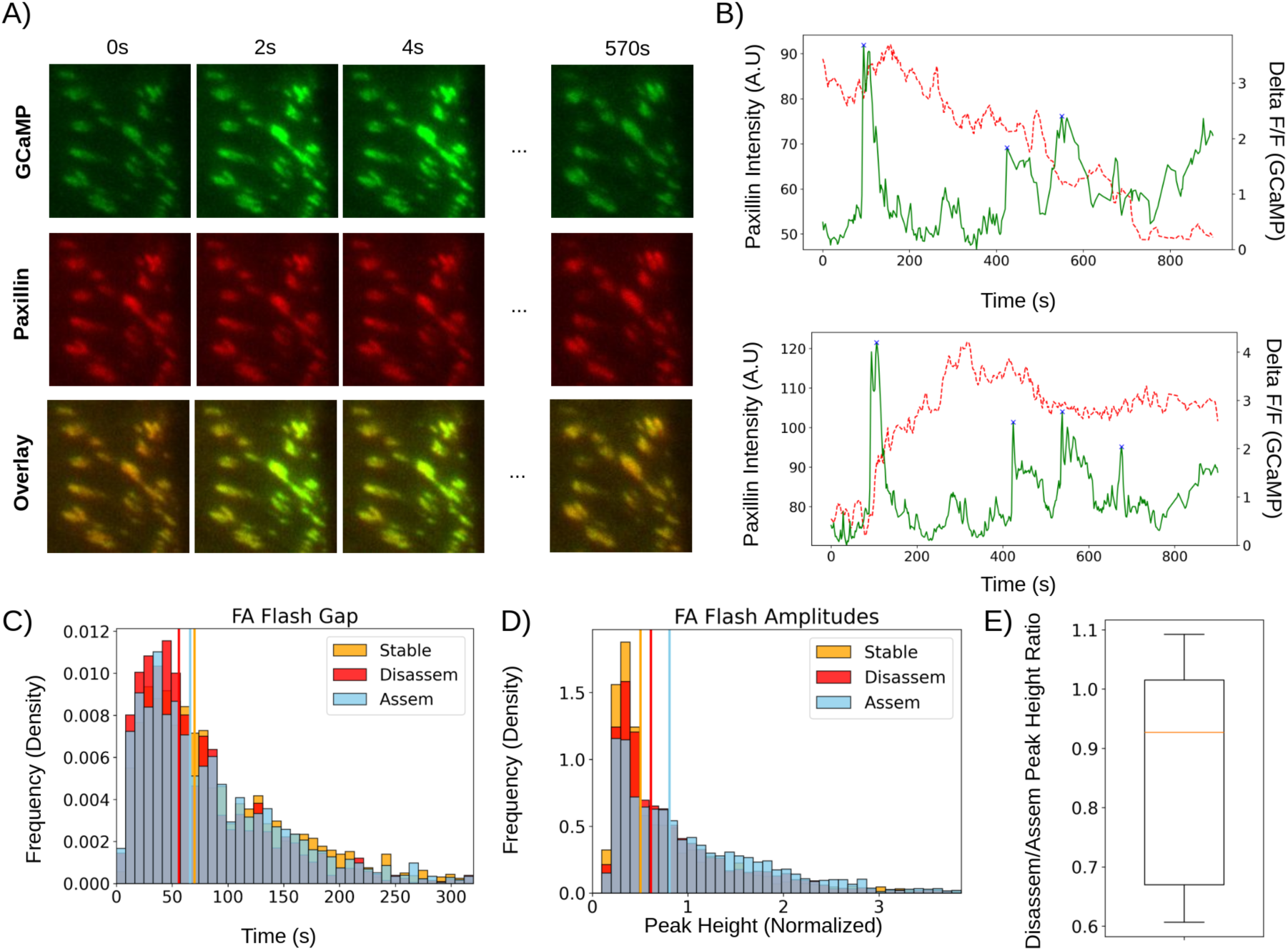
Calcium does not instigate FA disassembly in this system. (A) Representative examples of GCaMP6s and Paxillin Intensity at FAs over 570s. Images were acquired using live-cell TIRF imaging from U2OS cells. From top to bottom: GCaMP, Paxillin, Combined Channels. (B) Average GCaMP (dF/F) and Paxillin intensity over time. Green line: GCaMP, Red line: Paxillin. (C) Time between calcium flashes at disassembling, assembling and stable FAs. (D) Distribution of calcium flash amplitudes at disassembling, assembling and stable FAs. For (C-D), n=853 FAs across 13 cells. To improve clarity, histograms show all data up to the 99^th^ percentile. (E) Boxplot showing the average peak height of disassembling FAs divided by the average peak height of assembling FAs, done on a cell-cell basis; n=13 cells.

### Relaxation of SF contractility by retrograde actin flow drives mature FA disassembly

Given that calcium does not instigate mature FA disassembly, we considered the possibility that a mechanical stimulus, rather than signaling, drives FA disassembly. We reasoned that this putative instigator of FA disassembly likely disrupts some aspect of the signaling-contractility feedback loop and is probably coupled to cell movements like edge retraction that require FA disassembly. Based on these considerations, we hypothesized that retrograde actin flow may play a dual role in FA disassembly. This hypothesis hinges on the fact that many isoforms of myosin II, including nonmuscle myosin II (NMII), are load sensitive (22), and therefore exhibit reduced contractile force when exposed to a force in the direction of contraction (facilitating load). We reasoned that since retrograde actin flow tugs the FA in the same direction of the actomyosin contraction, this tugging is expected to facilitate and hence turn off the contractility, leading to FA disassembly.

To test this notion, we introduced in our model this retrograde actin flow-mediated relaxation effect on actomyosin contractility. While this addition did not notably change the model predictions on FA centripetal growth in Figure 2 and 3 (Supplemental Figure S2), reintroducing retrograde actin flow upon the mature FA exerts two effects on FAs: 1) it turns off stress fiber contractility, which reduces the traction at FAs and disengages catch bond reinforcement (Figure 5A), and 2) it carries unbound components downstream, which shifts the chemical balance towards detachment of anchored components (Figure 5B). Together, these two effects eventually cause the FA to fall out of the mechanical “goldilocks” zone (Figures 3G and 5C). This drives disengagement of anchored components ∼30s after the onset of the retrograde actin flow (Figure 5D) which results in FA disassembly (Figure 5E). Interestingly, the model predicts that FAs undergo “backsliding” as they disassemble, through a treadmilling process like that of nascent FA sliding (Figures 2C and 5D). This can potentially explain the phenomenon of “myosin-mediated backsliding” (45), which is known to occur at disassembling FAs but is not well understood.

**Figure 5:**
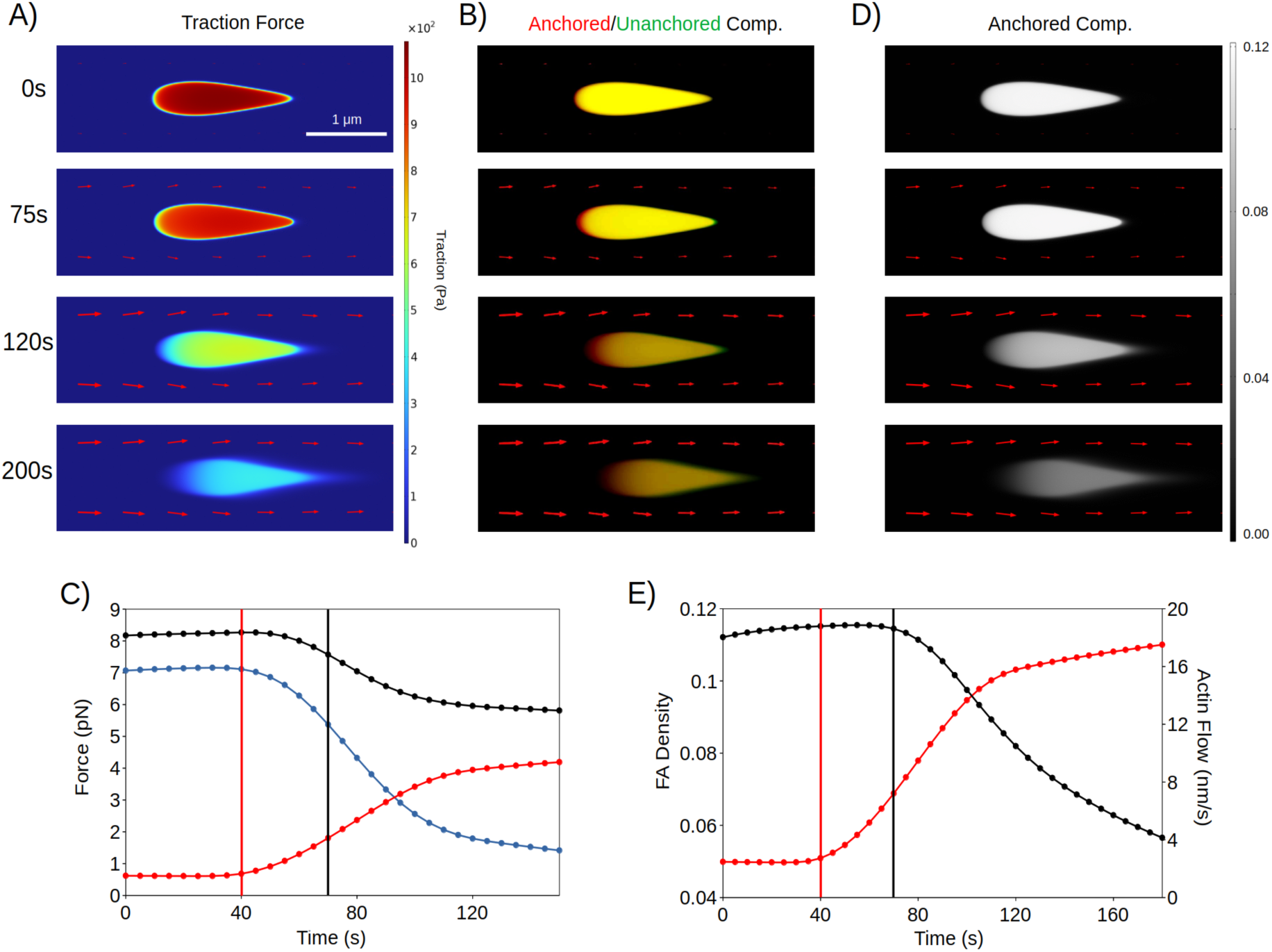
Mechanical interplay between actin flux and contractility drives FA disassembly. (A) FA traction over 200s, corresponding to timestamps in (C, E). (B) Distribution of anchored and unanchored components at disassembling FAs over 180s. (C) Force per anchored component (black, pN), force per component from stress fiber contractility (blue, pN) and force per component from retrograde actin flux (red, pN) over time. Vertical red line represents time of actin flux onset, while vertical black line represents start of FA disassembly, as seen in (E). (D) FA anchored density over 180s. (E) Average FA anchored component density and retrograde actin flux speed (nm/s) over time. Red and black plots represent actin flux speed and FA density. Vertical lines are as defined in (C).

Therefore, the model predicts that 1) the local retrograde actin flow is higher at disassembling adhesions compared to stable adhesions, 2) retrograde actin flow increases prior to the disassembly of mature FAs, and 3) retrograde actin flow is strongly anticorrelated with the size and density of FAs.

To test these predictions, we first experimentally measured the retrograde actin flow at disassembling FAs. We leveraged fluorescent speckle microscopy in combination with total internal reflection microscopy, which allowed us to image retrograde actin flow at the focal plane of FAs. We focused on the FAs at the retracting edge, which are expected to consist of mature FAs under significant retrograde actin flow. This revealed a substantial population of FAs undergoing disassembly in a manner like we predicted (Figure 6A). Guided by our model, we classified disassembling adhesions into flow-dependent and flow-independent modes based on a double gaussian fit (Figure 6B) and compared flow-dependent disassembling FAs to our model predictions. Notably, the flow-dependent population represented ∼ 92% of all disassembling FAs, suggesting it serves as the most common disassembly mode at retracting edges.

**Figure 6:**
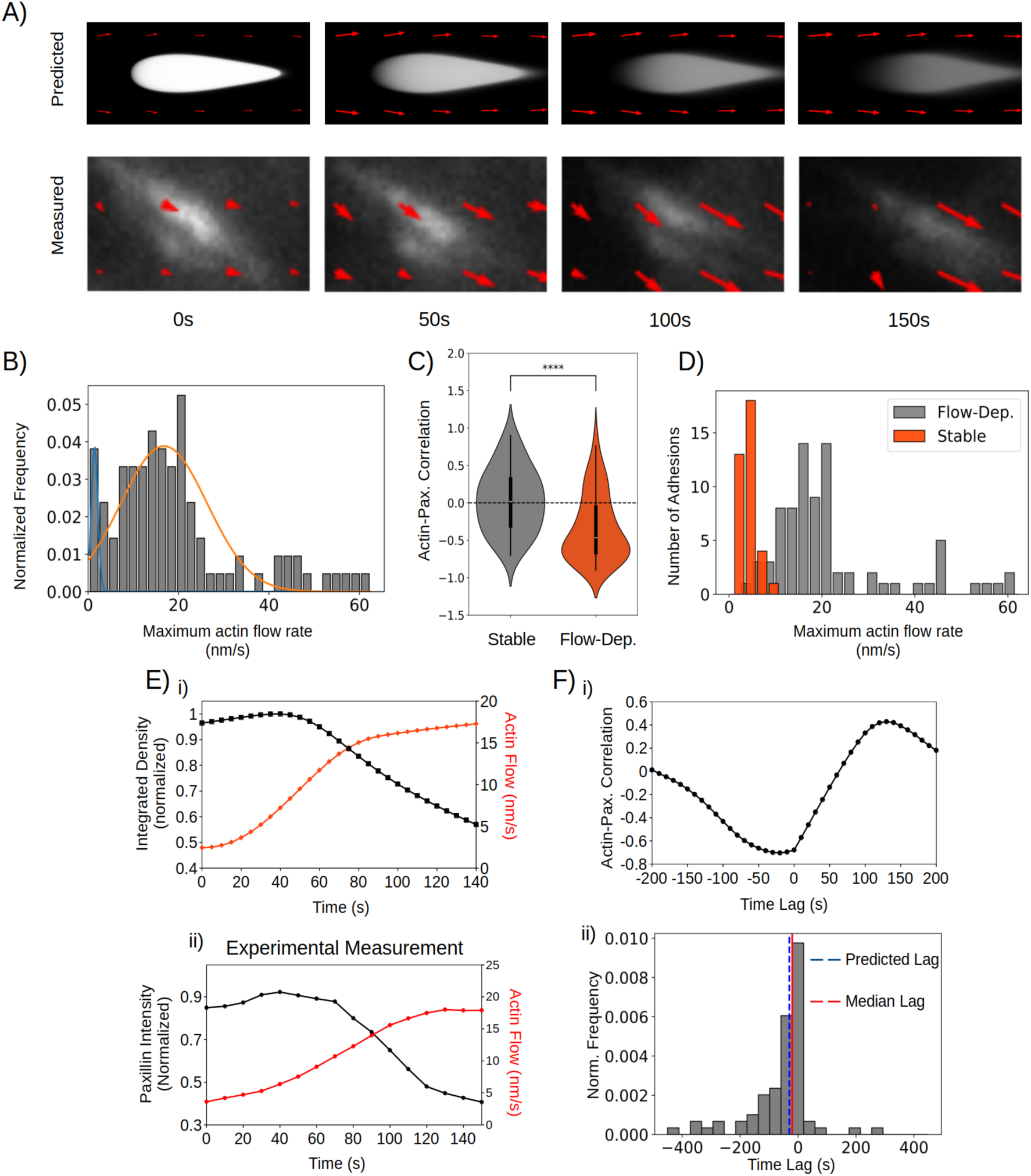
Actin flux is a key driver of FA disassembly at retracting edges. (A) Measured and simulated FA snapshots over 150s. (B) Gaussian mixture model of actin flux speed at disassembling FAs. Flow independent: µ = ∼1nm/s, weight = 0.08; flow-dependent: µ = ∼16nm/s, weight = 0.92. (C) Violin plot of Pearson Product-Moment Correlation Coefficient between actin flux and integrated FA density at stable (n=36, µ=0.019) and flow-dependent disassembling FAs (n=78, µ=-0.45). (D) Histogram of actin flow rates at stable and flow-dependent disassembling FAs. FAs undergoing flow-dependent disassembly are significantly more anticorrelated (stable: µ=0.019; flow-dep. disassembly: µ=-0.45; p < 1e-4, Mann-Whitney U-Test) and are under significantly higher flux (stable: µ=3.7nm/s, flow-dep. disassembly: µ=22nm/s; p < 1e-4, Mann-Whitney U-Test) compared to stable adhesions. (Ei) Predicted dynamics of actin flux and integrated FA density over time. Integrated FA density is in black, while actin flow speed is in red. (Eii) Experimentally measured integrated intensity and actin flow speed over time. Integrated intensity is in black, actin flow speed is in red. (Fi) Cross correlation of data in (Eii). (Fii) Distribution of time lags during flow-dependent disassembly (n=78). The median lag (20s) closely matches the predicted lag (30s).

To ensure retrograde actin flow is specific to disassembling FAs, we compared FAs undergoing flow-dependent disassembly with stable FAs and find that 1) paxillin is significantly more anticorrelated with retrograde actin flow for disassembling FAs than stable FAs (Figure 6C) and 2) that retrograde actin flow speeds at FAs undergoing flow-dependent disassembly are significantly higher compared to stable FAs (Figure 6D). Analysis of retrograde actin flow and FA intensity over time closely matches our theoretical prediction (Figure 6E) in timescale and dynamics. To further confirm retrograde actin flow drives FA disassembly, we considered the timing of actin flux with respect to FA disassembly. Cross-correlation analysis (Figure 6F(i)) between retrograde actin flux and integrated FA intensity revealed that the FA disassembly almost exclusively lags behind the increase in the local retrograde actin flow (∼94% of data has a time lag ≤ 0) with a median time lag of 20s (Figure 6F(ii)), closely matching our theoretical prediction. Taken together, our data strongly supports the model prediction that retrograde actin flow is the principal driver of mature FA disassembly.

## Discussion

In this work, we establish that without the need to invoke the additional pathways, cells modulate the spatial-temporal coordination between retrograde actin flow and SF to coherently drive the progression of the entire life cycle of FA. To ensure efficient force transmission, cell builds both FA and the associated SF-mediated actomyosin contractility as clutches that are in positive feedback with each other, strengthen upon mechanical resistance, and weaken upon relaxation. To reach this clutching state (*aka* mature FA), the retrograde actin flow promotes the centripetal growth of nascent FA in lamellipodium, the resulting FA-localized spatial gradient of PTK/PTP activities sets the stage for SF engagement, and the subsequent actomyosin contraction promotes the FA maturation (Figures 2 and 3). As the cell moves forward, the SF-engaging mature FA, while remaining stationary on ECM, “slides” backward relative to the frame of the cell, first to lamellum, where the retrograde actin flow diminishes, and eventually ends up at the rear of the cell. Now, to timely turn off the clutching and disassemble the mature FA at its rear, the cell adopts an ingenious solution: As the local cell edge retracts during cell migration, it naturally increases the local retrograde actin flow that tugs the clutching FA in the direction of the associated actomyosin contractility. It is this tugging that relaxes the contractile forces, turns off the clutching, and triggers the mature FA disassembly (Figures 5 and 6). Consequently, instead of being destroyed, the molecules that constitute this mature FA remain intact and can be readily reused for new FA assembly. This way, the FA-localized crosstalk between retrograde actin flow and SF seamlessly drives the entire life cycle of FA from the infancy, growth, maturation, and to turnover in coupling to cell migration.

We suggest that the above picture defines a principal mechanism underlying FA life cycle, because it efficiently utilizes the building materials for FA. However, what if mature FAs cannot be disassembled timely for whatever reasons, for instance, because the local retrograde actin flow is too small, or the FA-ECM binding is too strong (20)? In this case, cell may resort to additional pathways for FA disassembly. For instance, calcium influx is reported to activate calpain-mediated signaling axis that cleaves FA adaptor protein, talin, triggering the FA disassembly (24, 25). In our experimental conditions, while calcium influx does occur, it does not seem to instigate FA disassembly (Figure 4). In the future, it would be of great interest to determine what makes such difference and how the cell decision process is made to choose the different pathways of FA disassembly. The answers are expected to help us deepen our understanding of the organizational principles of FA-based cell migration.

Moreover, microtubule (MT) is reported to involve in FA’s life cycle (46–51). For instance, the plus ends of MTs sequester GEFH1, a Rho activator and, hence, regulate the contractility of the local actomyosins (51). Recent experiments demonstrated that when transiently approached by a MT, the FA undergoes a rapid GEFH1-mediated centripetal sliding and disassembly (45). It is proposed that the local release of GEFH1 from MT stimulates the local actomyosin contractility that physically disrupts the FA-ECM linkage, triggering the rapid FA disassembly (45). Our model can explain this interesting observation but from a different angle: while activating actomyosin contractility specifically in a stress fiber reinforces a FA, activating contractility in the surrounding actin network drives retrograde actin flow by pulling back the branched actin network (52). This local retrograde actin flow tugs the FA in the proximal direction, which turns off actomyosin contractility in the FA-localized stress fiber. The latter tips the chemical balance toward more FA anchored component converting to unbound components that drift with the retrograde actin flow downstream. This way, rather than the physical rupture of FA-ECM linkage, the retrograde actin flow drives the treadmilling of the FA, enroute to the FA disassembly (Figures 5B and 5D). This naturally explains the centripetal sliding of the disassembling FA, evidenced in experiments (45), which has not been accounted for. Additionally, the notion that retrograde actin flow and SF coordinate to shape the FA fate may help rationalize another set of interesting observations. While the local GEFH1 release from MT leads to FA disassembly as demonstrated in (45), the global MT disassembly stimulate FA formation and reinforcement (46). The puzzle here is: If the local increase in the contractility directly disrupts FAs as proposed in (45), then more GEFH1 will be released from the global MT disassembly; consequently, the contractility is expected to be stimulated more potently causing more FA disassembly, instead of the observed FA reinforcement. It could be that the global MT disassembly may impact more parameters than just the GEFH1 release-mediated contractility stimulation. Nevertheless, our model could provide a coherent explanation. According to our model phase diagram calculation, when the increase in the FA-engaging actomyosin contractility overrides that in the retrograde actin flow, the FA gets more strengthened (Supplemental Figure S3). Our future work will focus on dissecting whether and how the interplay between MT and actin cytoskeleton regulates the retrograde actin flow and actomyosin contractility in FA assembly/disassembly processes.

In summary, our work defines the coherent mechanistic basis of how a cell modulates retrograde actin flow and SF to drive the entire life cycle of FA and therefore balance the two opposing roles of FA in cell migration: *i.e.*, the efficient traction force transmission onto the ECM pulling the cell forward that requires the assembly of mature FAs clutching with the SF-mediated contractility, and the dissociation from the ECM conferring the cell rear retraction that entails the timely disassembly of mature FAs. This finding may shed light on the organizational principles that a cell streamlines the mechanochemical interplays between FA, cytoskeleton, and cell edge dynamics for an efficient migration.

## Acknowledgements

J. L. and S.P. designed and supervised the research; R. D. performed the computations of the theoretical model and image analysis; K. I. performed the experiments and image analysis. All authors analyzed the data and edited the manuscript. S. P. is supported by Canadian Institutes of Health Research (PJT-178272) and Natural Sciences and Engineering Research Council of Canada (RGPIN-2020-05881). J.L. is supported by Johns Hopkins University Startup Funds, Catalyst Awards, and National Institutes of Health RO1 GM148459.

---

**Figure S1:**
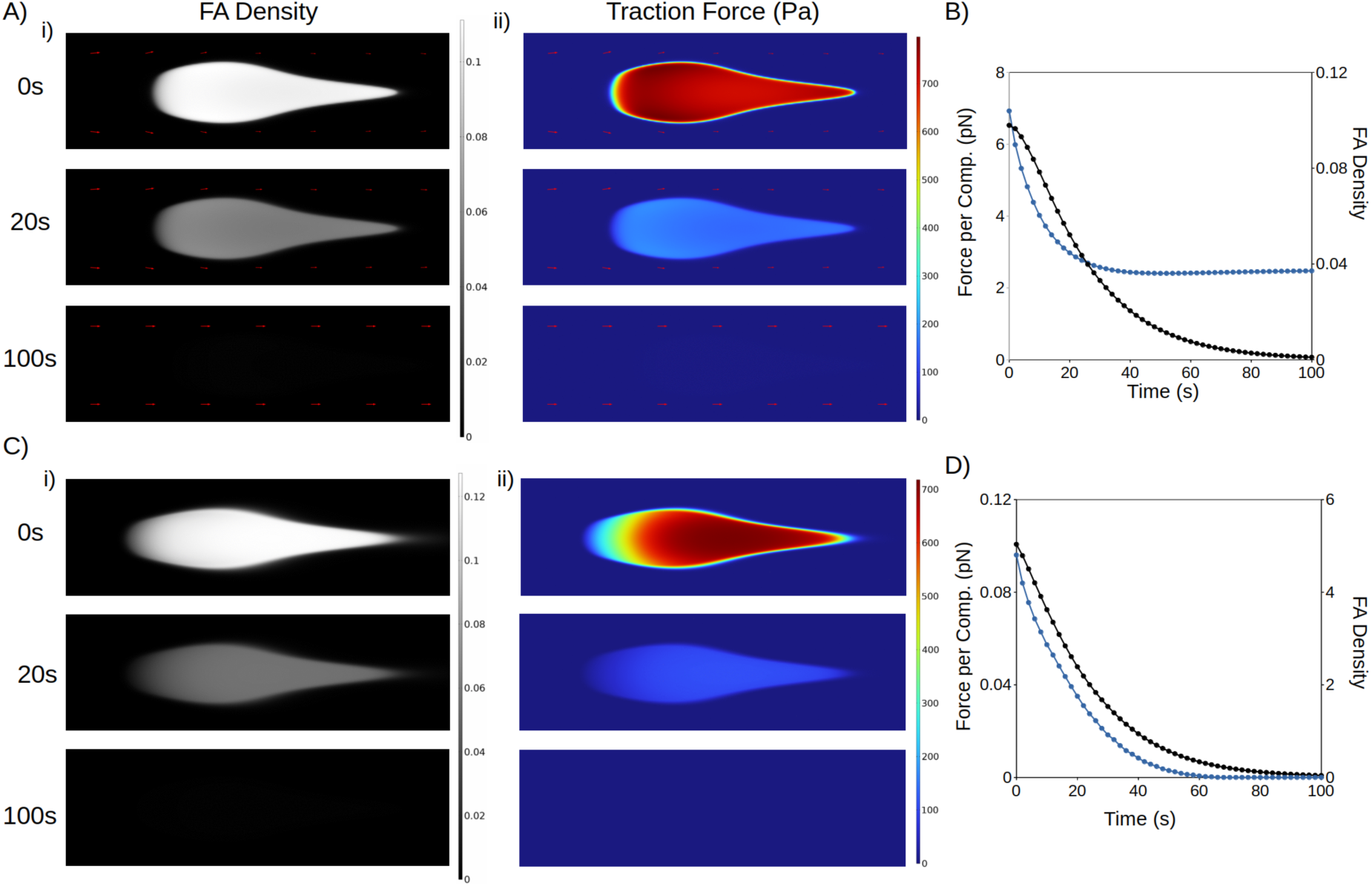
Substantial perturbations to contractility and the overlying actin network can drive FA disassembly. (Ai, ii) FA Density (Ai) and traction in Pascals (Aii) after reducing actomyosin contractility on-rate at the FA to zero. In the absence of actomyosin contractility, the rear of the FA becomes mechanically disengaged, leading to its disassembly. (B) Force per component and FA Density over 100s after setting contractility on-rate to zero. Force per component is in blue and FA density is in black. (Ci, ii) FA Density (Ci) and traction in Pascals (Cii) after reducing both retrograde actin flux and contractility on-rate to zero. With no source of mechanical engagement, catch-bonds become completely disengaged, leading to the disassembly of the entire FA. (D) Force per component and FA density over 100s after setting both flux and contractility on-rate to zero. Force per component is in blue and FA density is in black.

**Figure S2:**
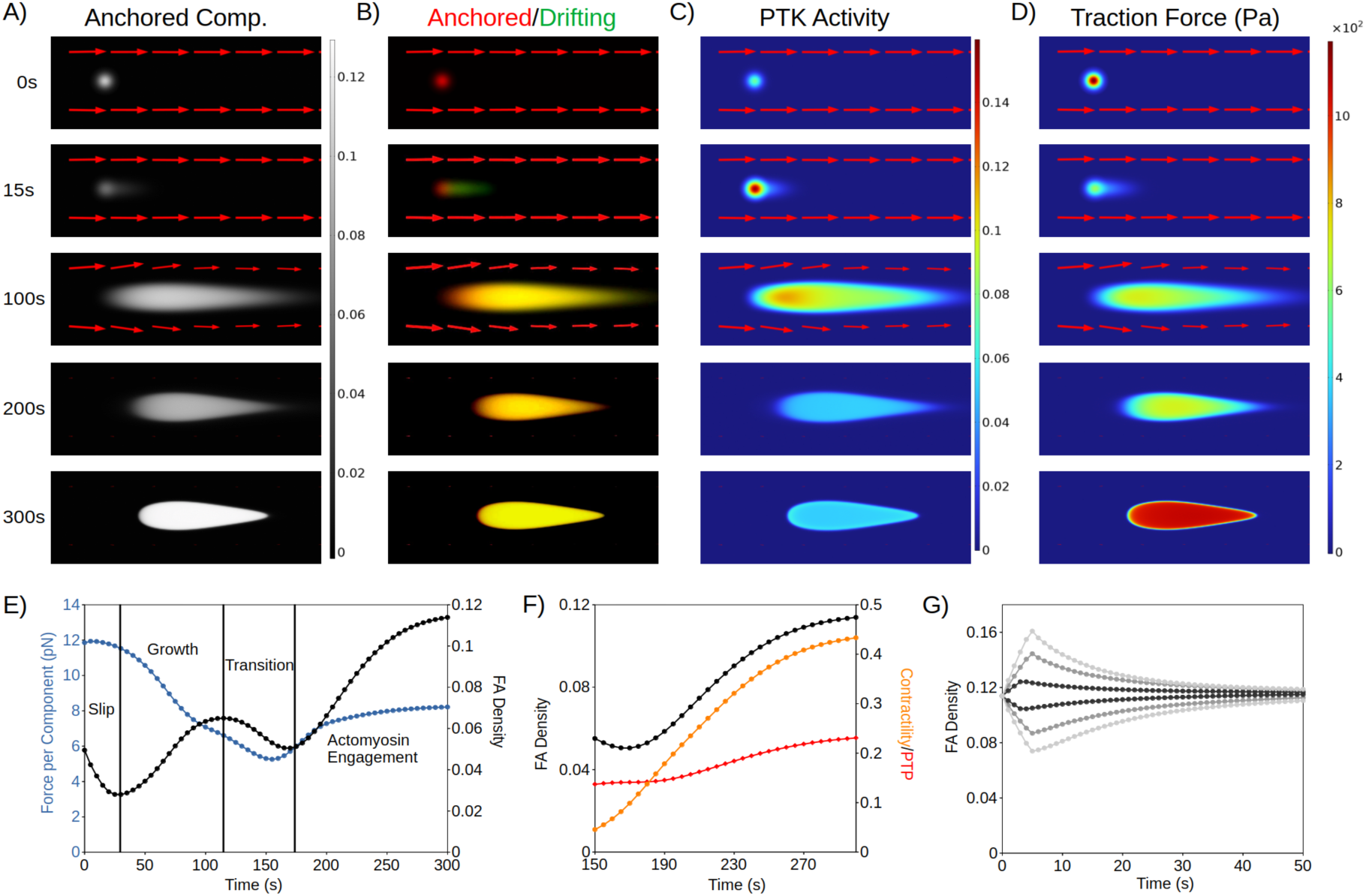
Inclusion of myosin load-dependence sustains key model predictions on FA assembly and stability. (A) Distribution of anchored FA components over 300s. Arrows indicate retrograde actin flux; length of arrows indicate flux magnitude. (B) Anchored and drifting FA components over 300s. Anchored components are in red, while drifting components are in green. (C) PTK Activity over 300s. The prediction that PTK is high during initial assembly and forms a gradient towards the distal end is preserved. (D). FA traction force over 300s. (E) FA Density and Force per Component (pN) over 300s. FA Density is in black, while Force per Component is in blue. The FA initially undergoes slipping upon engagement with actin, before forming a stronghold and growing. The FA then enters the lamella at 120s (modeled as a reduction of retrograde flux to 5nm/s), at which point the FA is sustained by actomyosin contractility. (F) Contractility, PTP activity and FA density from 150s to 300s after the start of the simulation. Contractility is in orange, PTP activity is in red, and FA density is in black. (G) FA Density in response to perturbations of different amplitude (0.05-0.15 1/s) exerted over 5s. The FA maintains stability in the face of small perturbations, as previously predicted.

**Figure S3:**
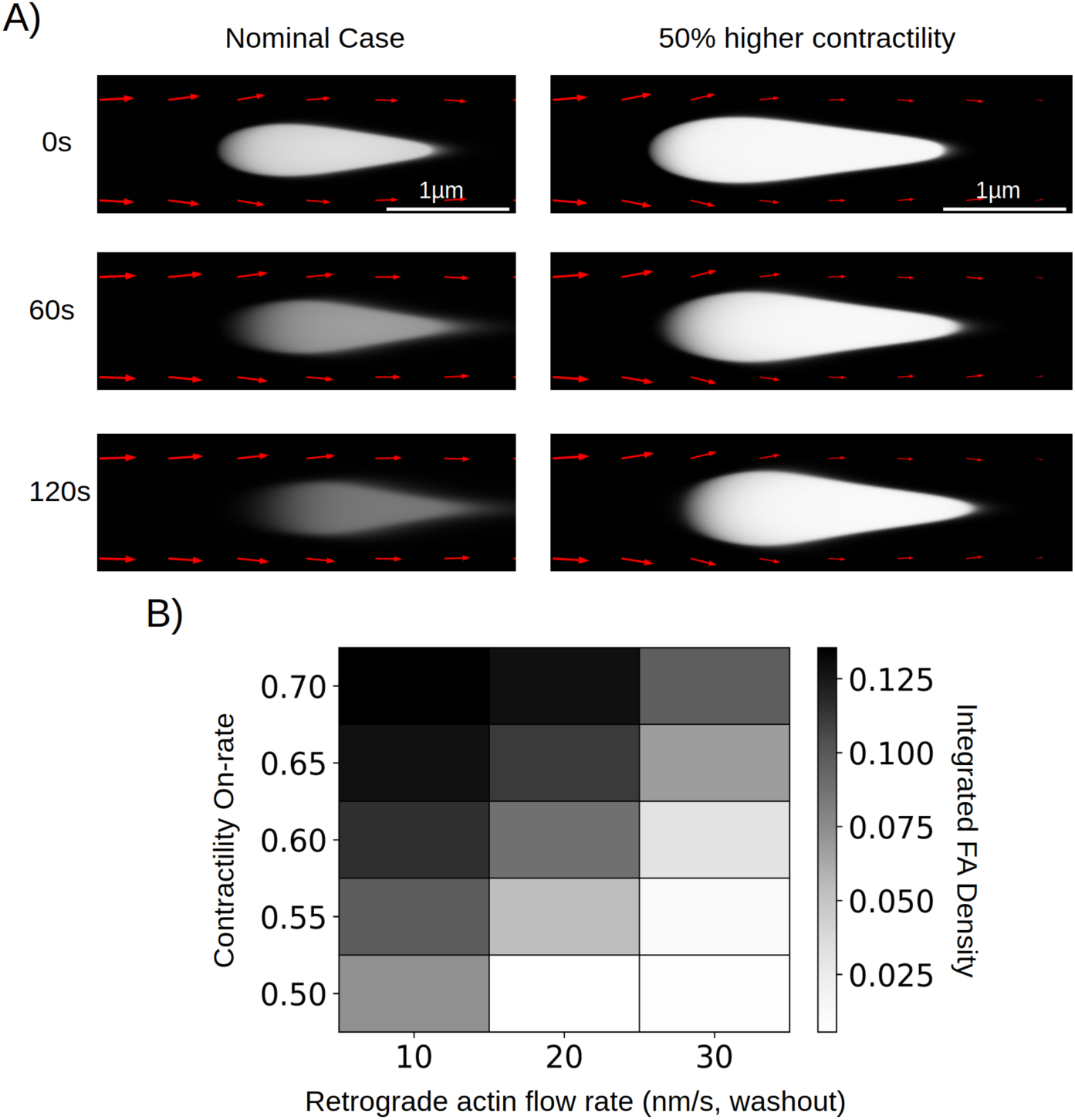
Increasing contractility in tandem with retrograde actin flow leads to FA growth. (A) Distribution of FA Density over 180s in the nominal case (parameters identical to Figure 5) and with 50% higher contractility (on-rate increased by 50%), mimicking the effect of global microtubule disassembly. (B) Phase plot of integrated FA density after 340s, at varying contractility on-rates (1/s) and retrograde actin flow rates after maturation in the lamella (nm/s).

## References

1. Alexander D. Bershadsky, Nathalie Q. Balaban, B. Geiger, Adhesion-Dependent Cell Mechanosensitivity. Annual Review of Cell and Developmental Biology 19, 677–695 (2003).

2. M. L. Gardel, I. C. Schneider, Y. Aratyn-Schaus, C. M. Waterman, Mechanical integration of actin and adhesion dynamics in cell migration. Annu Rev Cell Dev Biol 26, 315–333 (2010).

3. B. Geiger, J. P. Spatz, A. D. Bershadsky, Environmental sensing through focal adhesions. Nature Reviews Molecular Cell Biology 10, 21–33 (2009).

4. J. Z. Kechagia, J. Ivaska, P. Roca-Cusachs, Integrins as biomechanical sensors of the microenvironment. Nat Rev Mol Cell Biol 20, 457–473 (2019).

5. C. M. Lo, H. B. Wang, M. Dembo, Y. L. Wang, Cell movement is guided by the rigidity of the substrate. Biophys J 79, 144–152 (2000).

6. Sergey V. Plotnikov, Ana M. Pasapera, B. Sabass, Clare M. Waterman, Force Fluctuations within Focal Adhesions Mediate ECM-Rigidity Sensing to Guide Directed Cell Migration. Cell 151, 1513–1527 (2012).

7. Z. Wu, S. V. Plotnikov, A. Y. Moalim, C. M. Waterman, J. Liu, Two Distinct Actin Networks Mediate Traction Oscillations to Confer Focal Adhesion Mechanosensing. Biophysical Journal 112, 780–794 (2017).

8. L. A. Flanagan, Y. E. Ju, B. Marg, M. Osterfield, P. A. Janmey, Neurite branching on deformable substrates. Neuroreport 13, 2411–2415 (2002).

9. D. Koch, William J. Rosoff, J. Jiang, Herbert M. Geller, Jeffrey S. Urbach, Strength in the Periphery: Growth Cone Biomechanics and Substrate Rigidity Response in Peripheral and Central Nervous System Neurons. Biophysical Journal 102, 452–460 (2012).

10. A. J. Engler, S. Sen, H. L. Sweeney, D. E. Discher, Matrix Elasticity Directs Stem Cell Lineage Specification. Cell 126, 677–689 (2006).

11. W.-h. Guo, M. T. Frey, N. A. Burnham, Y.-l. Wang, Substrate Rigidity Regulates the Formation and Maintenance of Tissues. Biophysical Journal 90, 2213–2220 (2006).

12. M. J. Paszek et al., Tensional homeostasis and the malignant phenotype. Cancer Cell 8, 241–254 (2005).

13. T. A. Ulrich, E. M. de Juan Pardo, S. Kumar, The Mechanical Rigidity of the Extracellular Matrix Regulates the Structure, Motility, and Proliferation of Glioma Cells. Cancer Research 69, 4167–4174 (2009).

14. M. A. Wozniak, R. Desai, P. A. Solski, C. J. Der, P. J. Keely ROCK-generated contractility regulates breast epithelial cell differentiation in response to the physical properties of a three-dimensional collagen matrix. Journal of Cell Biology 163, 583–595 (2003).

15. C. Ballestrem, B. Hinz, B. A. Imhof, B. Wehrle-Haller Marching at the front and dragging behind : differential αVβ3-integrin turnover regulates focal adhesion behavior. Journal of Cell Biology 155, 1319–1332 (2001).

16. M. A. Digman, C. M. Brown, A. R. Horwitz, W. W. Mantulin, E. Gratton, Paxillin Dynamics Measured during Adhesion Assembly and Disassembly by Correlation Spectroscopy. Biophysical Journal 94, 2819–2831 (2008).

17. P. Kanchanawong et al., Nanoscale architecture of integrin-based cell adhesions. Nature 468, 580–584 (2010).

18. J. Liu et al., Talin determines the nanoscale architecture of focal adhesions. Proceedings of the National Academy of Sciences 112, E4864–E4873 (2015).

19. R. Kumari et al., Focal adhesions contain three specialized actin nanoscale layers. Nat Commun 15, 2547 (2024).

20. S. P. Palecek, A. Huttenlocher, A. F. Horwitz, D. A. Lauffenburger, Physical and biochemical regulation of integrin release during rear detachment of migrating cells. J Cell Sci 111 **(Pt** **7**), 929–940 (1998).

21. L. B. Case, C. M. Waterman, Integration of actin dynamics and cell adhesion by a three-dimensional, mechanosensitive molecular clutch. Nature Cell Biology 17, 955–963 (2015).

22. M. Kovács, K. Thirumurugan, P. J. Knight, J. R. Sellers, Load-dependent mechanism of nonmuscle myosin 2. Proceedings of the National Academy of Sciences 104, 9994–9999 (2007).

23. L. B. Case et al., Molecular mechanism of vinculin activation and nanoscale spatial organization in focal adhesions. Nat Cell Biol 17, 880–892 (2015).

24. S. J. Franco et al., Calpain-mediated proteolysis of talin regulates adhesion dynamics. Nat Cell Biol 6, 977–983 (2004).

25. K. T. Chan, D. A. Bennin, A. Huttenlocher, Regulation of adhesion dynamics by calpain-mediated proteolysis of focal adhesion kinase (FAK). J Biol Chem 285, 11418–11426 (2010).

26. C. Jurado, J. R. Haserick, J. Lee, Slipping or Gripping? Fluorescent Speckle Microscopy in Fish Keratocytes Reveals Two Different Mechanisms for Generating a Retrograde Flow of Actin. Molecular Biology of the Cell 16, 507–518 (2005).

27. K. Hu, L. Ji, K. T. Applegate, G. Danuser, C. M. Waterman-Storer, Differential transmission of actin motion within focal adhesions. Science 315, 111–115 (2007).

28. A. del Rio et al., Stretching Single Talin Rod Molecules Activates Vinculin Binding. Science 323, 638–641 (2009).

29. Y. Sawada et al., Force Sensing by Mechanical Extension of the Src Family Kinase Substrate p130Cas. Cell 127, 1015–1026 (2006).

30. K. Burridge, S. K. Sastry, J. L. Sallee, Regulation of Cell Adhesion by Protein-tyrosine Phosphatases: I. CELL-MATRIX ADHESION*. Journal of Biological Chemistry 281, 15593–15596 (2006).

31. N. Yokoyama, W. T. Miller, Protein phosphatase 2A interacts with the Src kinase substrate p130CAS. Oncogene 20, 6057–6065 (2001).

32. C. Guilluy, R. Garcia-Mata, K. Burridge, Rho protein crosstalk: another social network? Trends in Cell Biology 21, 718–726 (2011).

33. A. Tomar, D. D. Schlaepfer, Focal adhesion kinase: switching between GAPs and GEFs in the regulation of cell motility. Current Opinion in Cell Biology 21, 676–683 (2009).

34. P. Vallotton, S. L. Gupton, C. M. Waterman-Storer, G. Danuser, Simultaneous mapping of filamentous actin flow and turnover in migrating cells by quantitative fluorescent speckle microscopy. Proceedings of the National Academy of Sciences of the United States of America 101, 9660–9665 (2004).

35. C. K. Choi et al., Actin and α-actinin orchestrate the assembly and maturation of nascent adhesions in a myosin II motor-independent manner. Nature Cell Biology 10, 1039–1050 (2008).

36. X. Wang, T. Ha, Defining Single Molecular Forces Required to Activate Integrin and Notch Signaling. Science 340, 991–994 (2013).

37. M. Pittman et al., Membrane ruffling is a mechanosensor of extracellular fluid viscosity. Nature Physics 18, 1112–1121 (2022).

38. M. H. Jo et al., Molecular Nanomechanical Mapping of Histamine-Induced Smooth Muscle Cell Contraction and Shortening. ACS Nano 15, 11585–11596 (2021).

39. Y. Aratyn-Schaus, M. L. Gardel, Transient frictional slip between integrin and the ECM in focal adhesions under myosin II tension. Curr Biol 20, 1145–1153 (2010).

40. J. L. Guan, Role of focal adhesion kinase in integrin signaling. Int J Biochem Cell Biol 29, 1085–1096 (1997).

41. A. Y. Alexandrova et al., Comparative Dynamics of Retrograde Actin Flow and Focal Adhesions: Formation of Nascent Adhesions Triggers Transition from Fast to Slow Flow. PLOS ONE 3, e3234 (2008).

42. J. L. Tan et al., Cells lying on a bed of microneedles: An approach to isolate mechanical force. Proceedings of the National Academy of Sciences 100, 1484–1489 (2003).

43. X. Li, J. D. Combs, 3rd, K. Salaita, X. Shu, Polarized focal adhesion kinase activity within a focal adhesion during cell migration. Nat Chem Biol 19, 1458–1468 (2023).

44. R. Zaidel-Bar, R. Milo, Z. Kam, B. Geiger, A paxillin tyrosine phosphorylation switch regulates the assembly and form of cell-matrix adhesions. Journal of Cell Science 120, 137–148 (2007).

45. J. Aureille et al., Focal adhesions are controlled by microtubules through local contractility regulation. EMBO J 43, 2715–2732 (2024).

46. A. Bershadsky, A. Chausovsky, E. Becker, A. Lyubimova, B. Geiger, Involvement of microtubules in the control of adhesion-dependent signal transduction. Current Biology 6, 1279–1289 (1996).

47. I. Kaverina, K. Rottner, J. V. Small, Targeting, capture, and stabilization of microtubules at early focal adhesions. J Cell Biol 142, 181–190 (1998).

48. I. Kaverina, O. Krylyshkina, J. V. Small, Microtubule targeting of substrate contacts promotes their relaxation and dissociation. J Cell Biol 146, 1033–1044 (1999).

49. J. V. Small, B. Geiger, I. Kaverina, A. Bershadsky, How do microtubules guide migrating cells? Nat Rev Mol Cell Biol 3, 957–964 (2002).

50. E. J. Ezratty, M. A. Partridge, G. G. Gundersen, Microtubule-induced focal adhesion disassembly is mediated by dynamin and focal adhesion kinase. Nat Cell Biol 7, 581–590 (2005).

51. S. Stehbens, T. Wittmann, Targeting and transport: How microtubules control focal adhesion dynamics. Journal of Cell Biology 198, 481–489 (2012).

52. C. H. Lin, E. M. Espreafico, M. S. Mooseker, P. Forscher, Myosin drives retrograde F-actin flow in neuronal growth cones. Neuron 16, 769–782 (1996).

